# Broad application of plant protein tagging with ALFA tag for nanobody-based imaging and biochemical approaches

**DOI:** 10.1101/2025.03.19.644120

**Authors:** Julie Neveu, Julien Spielmann, Steven Fanara, Charlotte Delesalle, Sylvain Cantaloube, Grégory Vert

**Affiliations:** Plant Science Research Laboratory (LRSV), UMR5546 CNRS/University of Toulouse/Toulouse-INP, 24 chemin de Borde Rouge, 31320 Auzeville Tolosane, France; Light imaging Toulouse CBI (LITC), Centre de Biologie Integrative (CBI), FR3743 CNRS/University of Toulouse, 118 route de Narbonne, 31062 Toulouse CEDEX 9, France

**Author notes:** Co-first author (Julie Neveu, Julien Spielmann).

## Abstract

Epitope tags are broadly used for detecting, modifying or purifying proteins of interest, but their breadth of application is often rather poor. Recently, the rationally designed ALFA tag and its ALFA Nanobody complements the repertoire of epitope tags and emerged as a highly versatile epitope tag characterized in various animal models and that outperforms existing tags. Here we evaluated the ALFA tag/ALFA Nanobody technology in plants and showcase its application for *in planta* protein detection across multiple compartments or cellular structures, protein:protein interaction studies, protein purification, induced-proximity approaches, and super-resolution microcopy. Most importantly, we shed light on the edge of ALFA tagging technology for proteins difficult to tag for topological or functionals reasons. We provide the proof of concept of ALFA tag technology for the detection and functional analysis of the Arabidopsis IRT1 iron transporter. Overall, such versatile and validated toolbox of ALFA tag and ALFA Nanobody applications will serve as a resource of great interest for functional studies in plants.

## Introduction

Understanding the role of proteins of interest in plants requires precise tools to study their localization, interaction network, and regulation while maintaining their native structure and function. Fusion proteins with epitope tags or fluorescent tags are of central importance in protein science to facilitate their expression and purification, to immunopurify proteins and protein complexes, or to monitor their localization and dynamics (Tanz et al., 2013; Wang et al., 2016). Tag amino acid composition, size, affinity of corresponding tag-specific antibodies or recognition proteins greatly hamper the possibility of using a single tag for all cell biology and biochemical applications.

The recent successful application of small versatile tags recognized by nanobodies expressed independently in cells was reported in a variety of animal, yeast and bacterial models (Götzke et al., 2019; Traenkle et al., 2020; Xu et al., 2022). Among these, the ALFA tag was designed *de novo* to fulfill all criteria expected for a versatile tag for protein detection and purification (Götzke et al., 2019). Since its publication in animal cells, the ALFA tag technology system has been used for new applications and has been applied to other non-plant model organisms (Jedlitzke and Mootz, 2022; Westlund et al., 2023; Akhuli et al., 2024; Vicidomini et al., 2024). The use of nanobodies derived from the variable domain of camelid heavy-chain antibodies offers several advantages, including the possibility of being encoded in the genome of most model organisms, small size, high affinity, and their stability in cellular environments. These properties have been successfully harnessed for protein visualization (Keller et al., 2018), targeted degradation (Ju Shin et al., 2015), and functional studies (Shao et al., 2023) in various biological systems,(Schornack et al., 2009; Rocchetti et al., 2014). In plants, nanobodies have been used to track protein dynamics (Rocchetti et al., 2014), activate immune response (Wang et al., 2023), mislocalize protein (Schornack et al., 2009), force interactions (Rübsam et al., 2023) or induce degradation via the ubiquitin-proteasome pathway (Demidov et al., 2022). Nevertheless, most of these studies are based on endogenous epitopes or nanobodies recognizing the large GFP reporter, and fail to offer a global toolkit to study proteins of interest in plants.

In the present study, we show that the ALFA tag and corresponding ALFA Nanobody (NB) are a promising approach for advancing research in plant cell biology and plant protein biochemistry. We successfully expressed ALFA tag-based fusion proteins in stable transgenic plants to determine the subcellular localization of marker proteins in various subcellular compartments. We also demonstrate the benefit of ALFA tag to detect, immunopurify, image by super-resolution microscopy, or to use in induced-proximity approaches. Importantly, we extend our proof of concept to the study of the IRT1 transceptor (Dubeaux et al., 2018), which is notoriously hard to work with due to topology and difficulties to obtain functional fusions. Using the ALFA tag/ALFA NB technology, we confirmed the interaction of IRT1 with known partners by functional complementation, validated IRT1 metal-dependent endocytosis, and implemented high-end approaches previously inaccessible to IRT1 such as sptPALM or targeted degradation. The ALFA tag technology thus stands out as a tool of great interest to plant biologists and offers the unique opportunity to use a single epitope tag for a wide range of applications in plant biology.

## Results

### Expression of the ALFA NB-Fluorophore fusion in plants

The ALFA tag/NB-Fluorophore technology was developed as a two-component system where transgenic lines expressing ALFA NB-Fluorophore fusions are crossed with a POI-ALFA line or directly transformed with a POI-ALFA construct. To validate the ALFA tag strategy in plants, we first generated a variety of Arabidopsis stable transgenic lines ubiquitously expressing NB-Fluorophore fusions. To this purpose, we generated several binary vectors harboring the *UBIQUITIN10* promoter (Norris et al., 1993) (*UBI10*), the *ALFA NB* codon optimized for Arabidopsis, a linker, and various fluorescent proteins (FP) (Fig. 1A). The FP used in this study are mCitrine (mCit) (Zacharias et al., 2002), tag-BFP (Subach et al., 2008), mScarlet (mSca) (Bindels et al., 2017), mEos 3.2 (mEos) (Zhang et al., 2012) and the photoswitchable DRONPA (Ando et al., 2004) (Fig. S1). We favored the use of the *UBI10* promoter rather than *CaMV35S* for its ability to yield mild constitutive expression levels and for the absence of silencing that is often observed with *CaMV35S*. *Arabidopsis* plants were transformed with these vectors and mono-insertional homozygous T3 lines were selected before undergoing phenotypic analyses. No interference with normal plant growth and development could be observed, indicating that *UBI10*-driven ALFA NB-FP fusions are not toxic to plant cells (Fig. 1 B-D and Fig. S1). Confocal microscopy observations revealed a strong fluorescent signal in root tips, differentiated root cells including root hairs, hypocotyls, mesophyll cells and guard cells. In all cases, the ALFA NB-FP fusion proteins localized in the cytoplasm and the nucleus (Fig. 1 E-J). Altogether, these results suggested that ALFA NB-Fluorophore is suitable for functional studies *in planta*.

**Figure 1.**
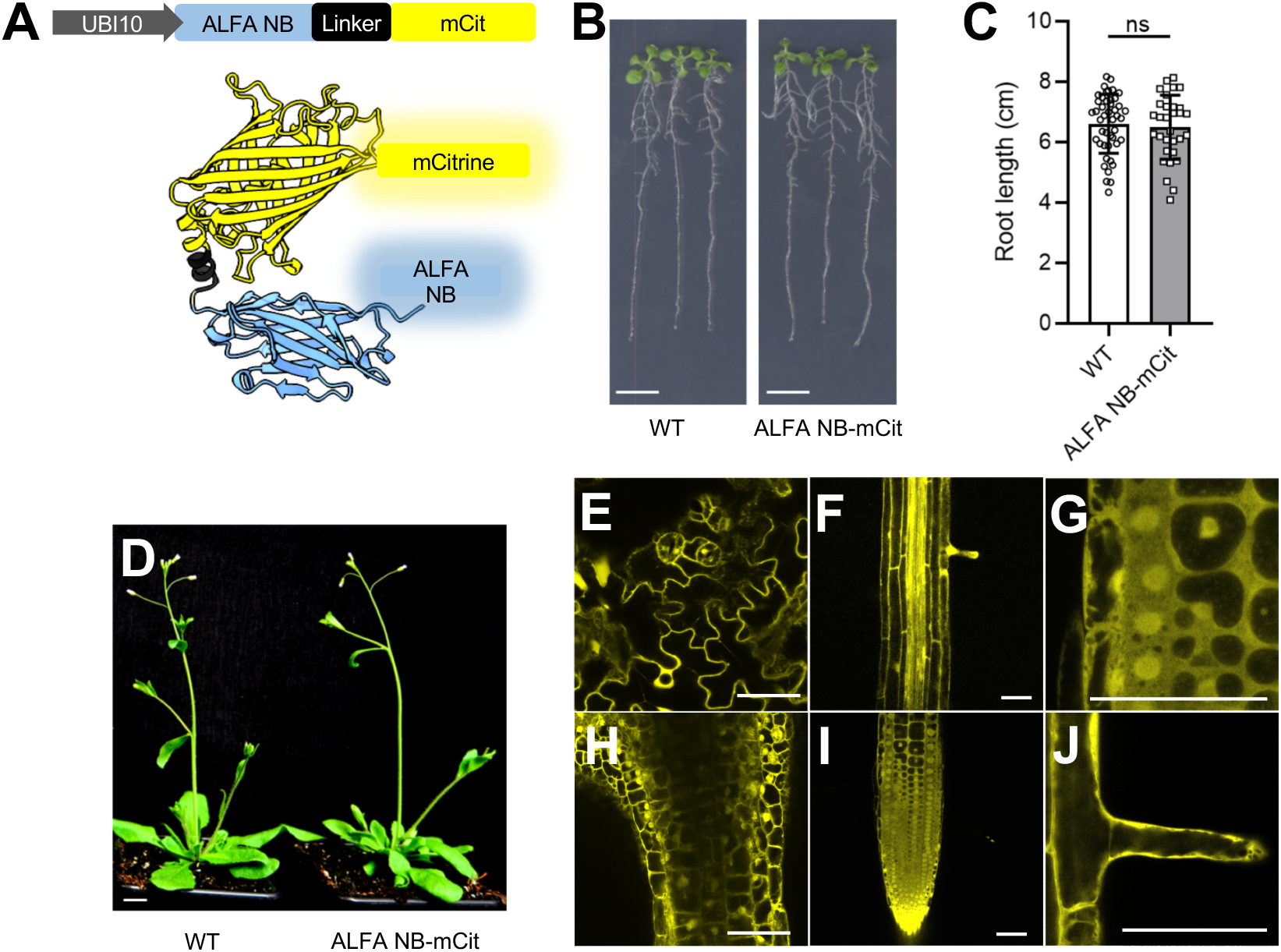
Expression of ALFA Nanobody-fluorescent protein fusions in plants. (A) Schematic illustration of the ALFA Nanobody (ALFA NB) fusion to mCitrine (mCit) expressed in *A. thaliana*. (B) Comparison of 14-day-old Arabidopsis wild-type (WT, Col-0) and transgenic plants expressing ALFA NB-mCit fusion. (C) Statistical analysis of root length of 14-day-old Arabidopsis seedlings grown on half-strength MS medium. Experiments were carried out in triplicates. Error bars represent standard error (n=30). No statistical difference was observed (Student’s t-test; ns, not significant). (D) Representative phenotypes of wild-type and ALFA NB-mCit plants. Scale bar 1cm. (E-J) Confocal microscopy images of Arabidopsis plants expressing ALFA NB fused to mCit in the leaves (E), roots (F-G), hypocotyls (H), root tips (I) and root hairs (J). Scale bar 50µm.

### Protein localization using the ALFA NB-FP technology

To demonstrate the ability of ALFA NB-FPs to serve as a tool to indirectly monitor the subcellular localization of a protein of interest, we first examined whether ALFA NB-FP could detect established markers of different subcellular compartments or cellular entities fused to the ALFA tag. Plants expressing the ALFA NB-mCit fusion protein were transformed with a construct allowing the expression of the Lti6b-ALFA fusion. In contrast to plants expressing ALFA NB-mCit only that show nucleocytoplasmic fluorescence (Fig. 1E-F), co-expression of Lti6b-ALFA resulted in plasma membrane localization of the ALFA NB-mCit (Fig. 2A). No fluorescence was observed in the cytoplasm or the nucleoplasm, indicating that Lti6b-ALFA recruited most of the NB reporter to the cell surface. A few cells however harbored tonoplastic localization of the ALFA NB-mCit (Fig. 2A). Considering that Lti6b is a plasma membrane protein, it undergoes endocytosis and intracellular sorting into multivesicular bodies (MVBs) on its way to the vacuole. Lti6b-ALFA bound to ALFA NB-mCit fails to be properly sorted into MVBs and likely ends up on the tonoplast, as already reported for missorted proteins in MVB (Cai et al., 2014).

**Figure 2.**
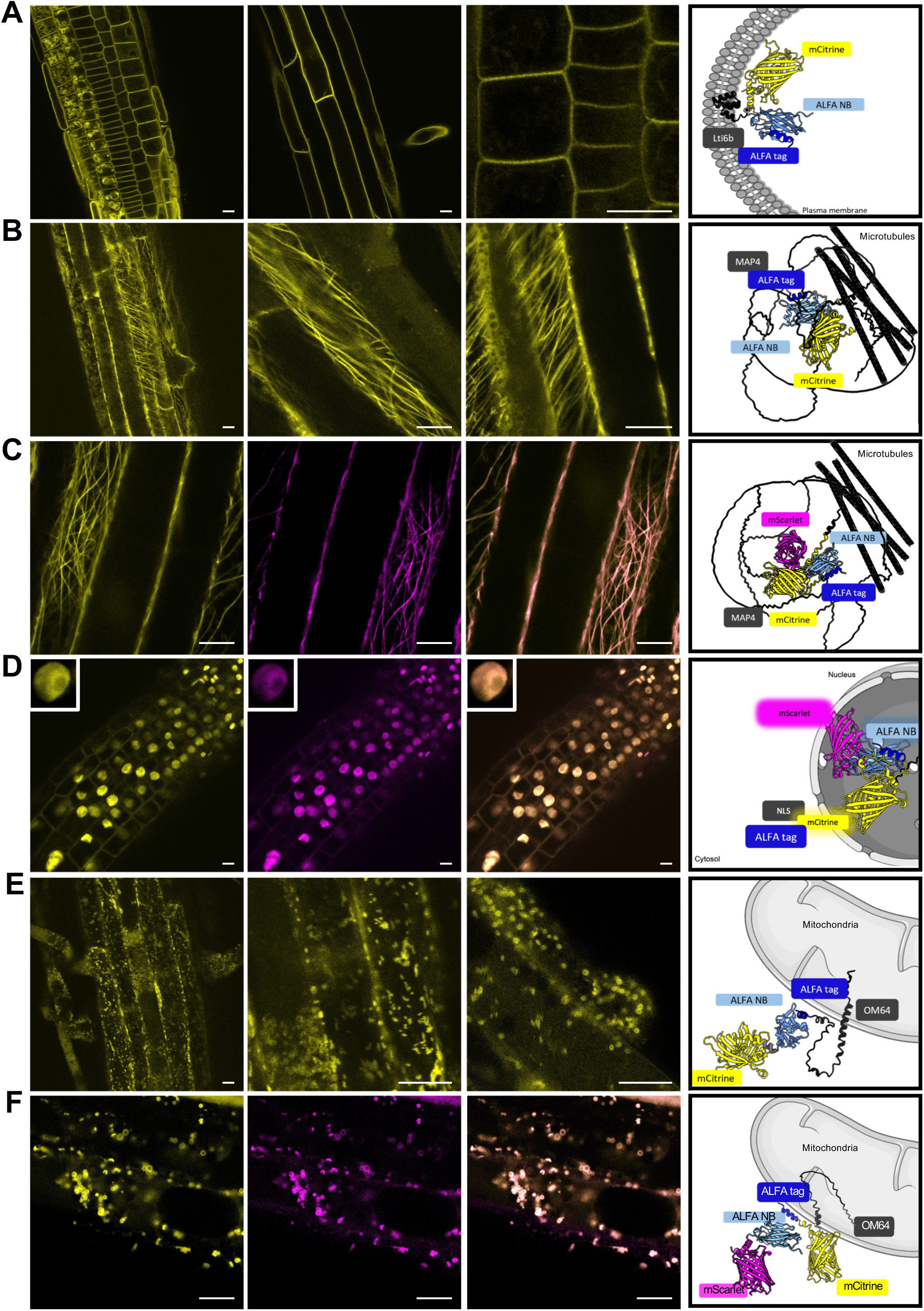
Detection of ALFA-tagged proteins by ALFA NB fused to fluorescent proteins. (A-E) Schematic representations and representative confocal microscopy images of Arabidopsis plants co-expressing Lti6b-ALFA and ALFA NB fused to mCit (A), MAP4-ALFA and ALFA NB fused to mCit (B), MAP4-mCit-ALFA and ALFA NB fused to mSca (C), NLS-mCit-ALFA and ALFA-NB fused to mSca (D), OM64-ALFA and ALFA NB fused to mCit (E), N and OM64-mCit-ALFA and ALFA NB fused to mSca (F). Scale bars 10µm. Illustrations were created using BioRender.com.

We then used the microtubule-binding domain of MAP4 (Marc et al., 1998) to assess whether ALFA NB-FPs can be used to study microtubule-associated proteins. Plants co-expressing ALFA NB-mCit and the MAP4-ALFA fusion showed a filamentous pattern that resembles typical microtubule staining (Fig. 2B). To ascertain that the observed pattern represented microtubules, plants were exposed to the microtubule-depolymerizing drug oryzalin (Morejohn et al., 1987). Oryzalin resulted in the loss of filamentous pattern (Fig. S2), suggesting that ALFA NB-mCit detected microtubule-associated ALFA-tagged MAP4. This is further substantiated by the fact that ALFA-tagged MAP4 fused to mCit clearly colocalized with ALFA NB-mSca (Fig. 2C).

We next investigated if such a reporter could be harnessed to study the localization of proteins not localized in the cytoplasm. When ALFA NB-mSca was co-expressed with a fusion between the Nuclear Localization Signal (NLS) sequence from the SV40 virus Large T-antigen (Collas et al., 1996), the ALFA tag and mCit, ALFA-mSca no longer displayed cytoplasmic fluorescence and perfectly matched with the NLS-driven nuclear localization of mCit (Fig. 2D). This confirms that the ALFA NB-FP enters the nucleus, consistent with its size being below the size exclusion limit of 40-50kDa, and is able to detect ALFA-tagged nuclear proteins. Similarly, the ALFA NB-mCit was also able to recognize the OM64 outer mitochondrial membrane protein (Murcha et al., 2014), which is a widely used mitochondrial marker, if harboring the ALFA tag towards the cytoplasm (Fig. 2E). Additionally, plants co-expressing ALFA-tagged OM64 fused to mCit and ALFA NB-mSca exhibited a perfect colocalization between mCit and mSca (Fig. 2F). In contrast the ALFA NB-mSca failed to detect the IM inner mitochondrial addressing signal fused to ALFA tag and mCit (Fig. S3A). Indeed, although the ALFA-IM-mCit fusion displayed a typical punctate pattern corresponding to mitochondria, the ALFA NB-mSca remained mostly diffuse in the cytoplasm and did not reach the inner mitochondrial membrane. The ALFA NB also proved to be ineffective to reach the lumen from the endoplasmic reticulum where the HDEL-mSca fused to the ALFA tag localizes (Fig. S3B).

Altogether, our observations argue that the ALFA NB fused to a fluorescent protein is suitable to monitor the subcellular localization of proteins in plants.

### Tagging of the IRT1 root iron transporter with ALFA tag

The bipartite ALFA NB/ALFA tag reporter system emerges as an important tool for imaging proteins that are hard to tag without impairing their function in plant cells. This is notably the case of many hydrophobic multispan membrane proteins for which N- and C-termini i) cannot be tagged without altering functionality, or ii) are exposed to acidic pH that impacts fluorescent proteins. A perfect example is the IRT1 root metal transporter from the model plant *Arabidopsis thaliana* for which only the insertion of a fluorescent protein in an internal loop facing the extracellular space allows the monitoring of the localization of a functional IRT1 protein while maintaining iron transport activity (Dubeaux et al., 2018). To showcase the benefit of the ALFA NB/ALFA tag reporter system, we set out to generate functional fusions of IRT1 with the ALFA tag. To this purpose, we generated different IRT1 fusions with the ALFA tag and expressed them in the *irt1* CRISPR knock-out mutant (*irt1_crispr_*) using complementation of lead chlorosis as a proxy for functionality. IRT1-ALFA fusions were designed by inserting the ALFA tag into different loops of the IRT1 protein (Fig. 3A). The first insertion (#1) was placed in the first extracellular loop, between residues P41 and C42, where a previously reported mCit was inserted and yielded functional IRT1 fusion (Dubeaux et al., 2018). Such *irt1crispr*/IRT1-ALFA fusion was functional as evidenced by the complementation of the severe chlorosis and growth defect of *irt1* loss of function mutant, serving as complementation control for the other fusions (Fig. 3B). The other IRT1-ALFA fusions were designed with the ALFA tag inserted into cytosolic loops to allow IRT1 to be targeted *in vivo* by the ALFA NB. Three insertion positions were tested, between S73 and R74 (#2), between T148 and S149 (#3) and between S322 and I323 (#4). Of these different fusions, only those at positions 1 and 3 were able to complement the *irt1_crispr_* mutant (Fig. 3B) and thus considered functional.

**Figure 3.**
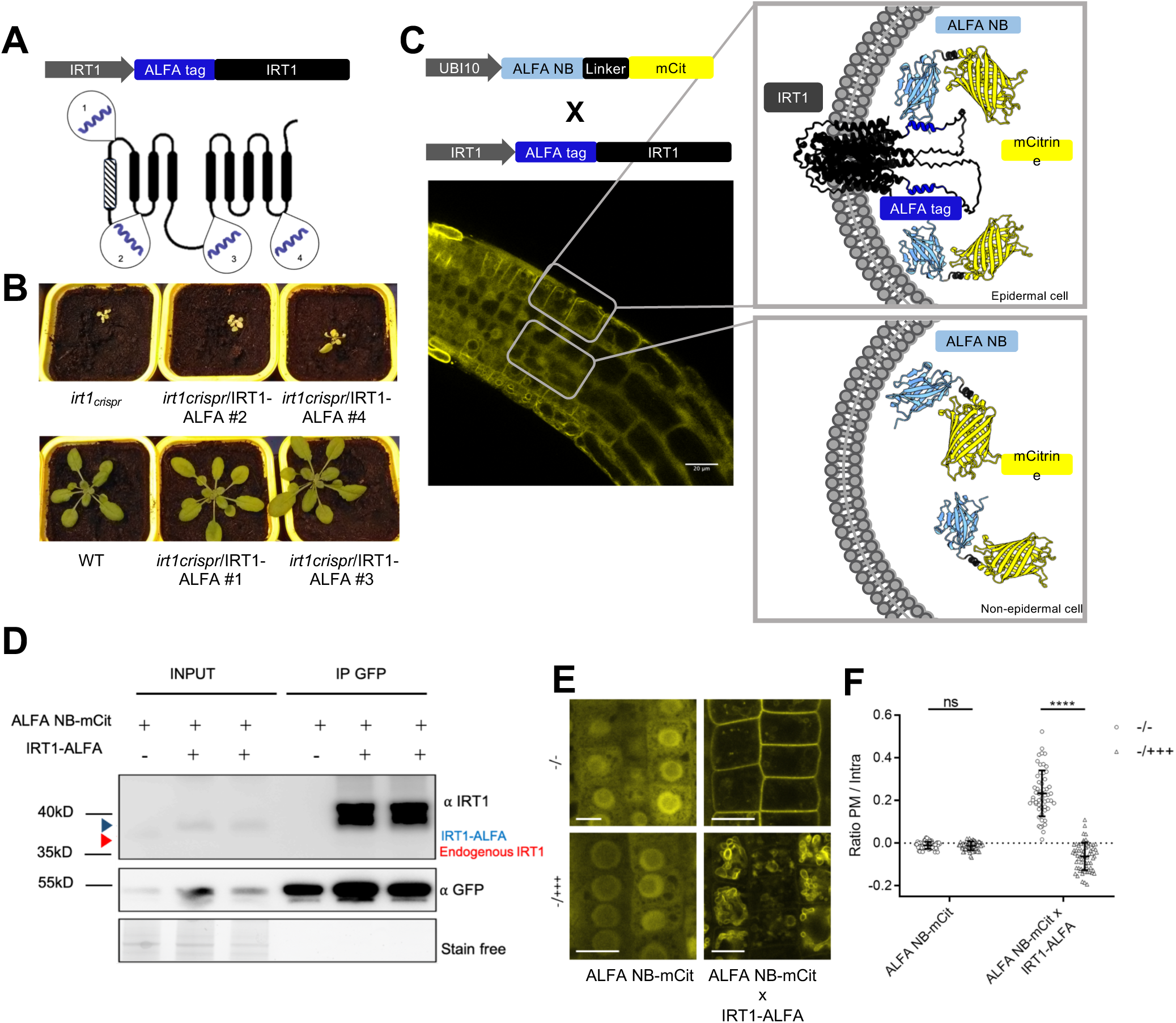
Proof of concept for ALFA technology applied to IRT1 imaging and biochemical detection. (A) Schematic representation of the ALFA tag insertion sites in IRT1 protein tested. (B) Complementation of *irt1* CRISPR mutant phenotype with pIRT1::IRT1-ALFA constructs, upper left to lower right, *irt1_crispr_*, *irt1_crispr_*/pIRT1::IRT1-ALFA#2, *irt1_crispr_*/pIRT1::IRT1-ALFA#4, WT, *irt1_crispr_*/pIRT1::IRT1-ALFA#1 and *irt1_crispr_*/pIRT1::IRT1-ALFA#3. (C) Confocal microscopy images of root from IRT1-ALFA plants expressing ALFA NB fused to mCit and schematic representation of epidermal cells expressing IRT1-ALFA and non-epidermal cells with no expression of IRT1-ALFA (BioRender). (D) Immunoprecipitation (IP) of IRT1-ALFA with ALFA NB-mCit. IP was performed using anti-GFP antibodies on protein extracts from 10-day-old seedlings, grown on half-strength MS without Fe, expressing ALFA NB-mCit alone or co-expressing ALFA NB-mCit and IRT1-ALFA. Immunoprecipitated proteins were subjected to immunoblotting with anti-GFP and anti-IRT1 antibodies. (E) Confocal microscope analysis of root epidermal cells from plants expressing ALFA NB-mCit alone or co-expressing ALFA NB-mCit and IRT1-ALFA. Plants were grown without Fe and non-Fe metals (-/-) and then liquid-treated for 24 h with either the same medium (-/-), or with an excess of non-Fe metals (-/+++). Scale bars, 10 µm. (F) Quantification of plasma membrane to intracellular signal ratio for plants expressing ALFA NB-mCit alone or co-expressing ALFA NB-mCit and IRT1-ALFA shown in (E). Error bars represent standard deviation and asterisks indicate statistically significant differences [two-way ANOVA, Sidak post hoc test; ****P < 0.0001; ns, not significant].

### ALFA tag-based imaging and biochemical detection of IRT1

Complemented *irt1_crispr_*/IRT1-ALFA#3 was crossed with plants expressing ALFA NB-mCit to monitor the subcellular localization of IRT1-ALFA fusions in root epidermal cells where endogenous IRT1 is found (Vert et al., 2002). IRT1-ALFA, detected through genetically encoded ALFA NB-mCit, showed plasma membrane localization in epidermal cells, as expected. IRT1-ALFA also localized to the tonoplast, highlighting again the possible interference of ALFA NB-mCit bound to IRT1-ALFA in late endosomal sorting processes (Fig. 3C). Since the ALFA NB-mCit reporter is expressed under the control of a constitutive promoter, fluorescence was also observed in non-epidermal cell layers where IRT1 is not expressed and showed a diffuse cytosolic localization (Fig. 3C). This demonstrates the targeting specificity of the ALFA NB. To evaluate the use of the ALFA-tag system for biochemical approaches and more specifically the study of protein regulation, we first validated the possibility of co-purifying the protein fused to the ALFA tag with the ALFA NB. We co-immunoprecipitated IRT1-ALFA from *irt1_crispr_*/IRT1-ALFA/ALFA NB-mCit plants using antibodies directed against GFP and capable of recognizing mCit (Fig. 3D). ALFA NB-mCit plants in wild-type background were used as control. Input fractions showed endogenous IRT1 protein detected with anti-IRT1 antibodies only for ALFA NB-mCit plants, consistent with *irt1crispr*/IRT1-ALFA/ALFA NB-mCit being devoid of endogenous IRT1. ALFA NB-mCit was detected using anti-GFP antibodies in all lanes. IRT1-ALFA was successfully co-immunoprecipitated from IRT1-ALFA-expressing plants with ALFA NB-mCit, as evidenced by signal observed at 40kDa, while no signal could be observed for ALFA NB-mCit alone (Fig. 3D).

We previously reported that IRT1 is endocytosed and targeted to the vacuole upon non-iron metal excess(Dubeaux et al., 2018; Spielmann et al., 2022). We therefore wondered whether the ALFA tag system could be used to visualize the change in distribution of IRT1 in response to metals. Plants expressing ALFA NB-mCit only (control) or *irt1crispr*/IRT1-ALFA/ALFA NB-mCit were subjected to different non-iron metal deficiency or excess and the localization of ALFA NB-mCit monitored by confocal microscopy as a proxy for IRT1-ALFA. ALFA NB-mCit localized to the cytoplasm and nucleus irrespective of non-iron metals (Fig. 3E). However, when co-expressed with IRT1-ALFA in the absence of non-iron metals, ALFA NB-mCit nicely localized to the plasma membrane where IRT1 is responsible for iron uptake. Metal excess radically changed IRT1-ALFA localization detected through ALFA NB-mCit, as highlighted by the removal from the cell surface and targeting to the vacuole. As previously observed, ALFA NB-mCit interfered with vacuolar delivery and mostly yielded accumulation to the tonoplast. Quantifying the plasma membrane to intracellular fluorescence ratio supports the strong impact of metal excess on IRT1 endocytosis (Fig. 3F).

### ALFA tag-based fluorescence complementation for IRT1 protein:protein interaction studies

ALFA tag-based fusion proteins also offer the possibility to perform fluorescence complementation approaches to monitor and localize protein:protein interaction for proteins of interest with extracellular N- and C-termini and/or that cannot be tagged without impairing activity. To illustrate this, we have once again taken advantage of the IRT1. Both the functional IRT1-mCit fusion, where mCit is inserted into the first extracellular loop of IRT1, and the functional IRT1-ALFA fusion 3 carrying the ALFA tag inserted in the large cytosolic loop of IRT1 display plasma membrane localization when transiently expressed in *Nicotiana benthamiana* leaves (Fig. 4A). Bifunctional fluorescence complementation experiments using N-terminally tagged IRT1 with half of mCitN and proteins known to interact with IRT1, C-terminally tagged with mCitC, such as FRO2 and AHA2 (Martín-Barranco et al., 2020) revealed mCit reconstitution both intracellularly and at the plasma membrane (Fig. 4B). This is likely explained by the cleavage of the predicted signal peptide in N-ter of IRT1 that releases mCitN and results in non-specific interaction with mCitC-tagged IRT1 partners. Consistently, co-expression of a mCitN-IRT1 fusion protein with mCitC led to a non-specific fluorescent signal in the nucleus. To circumvent these issues, we took advantage of the ALFA tag/ALFA NB technology and designed a tripartite functional fluorescence complementation assay (TriFC) using the functional IRT1-ALFA, with the ALFA tag being found in an intracellular loop, the ALFA NB-mCitN fusion, and the mCitC-tagged IRT1 partners. Co-expression of TriFC components revealed a clear plasma membrane mCit reconstitution with AHA2-mCitC and FRO2-mCitC (Martín-Barranco et al., 2020), but not with ALFA-Lti6b negative control (Fig. 4C, D). These results expand the possibility of using fluorescence complementation approaches to proteins difficult to tag and/or with unfavorable topologies.

**Figure 4.**
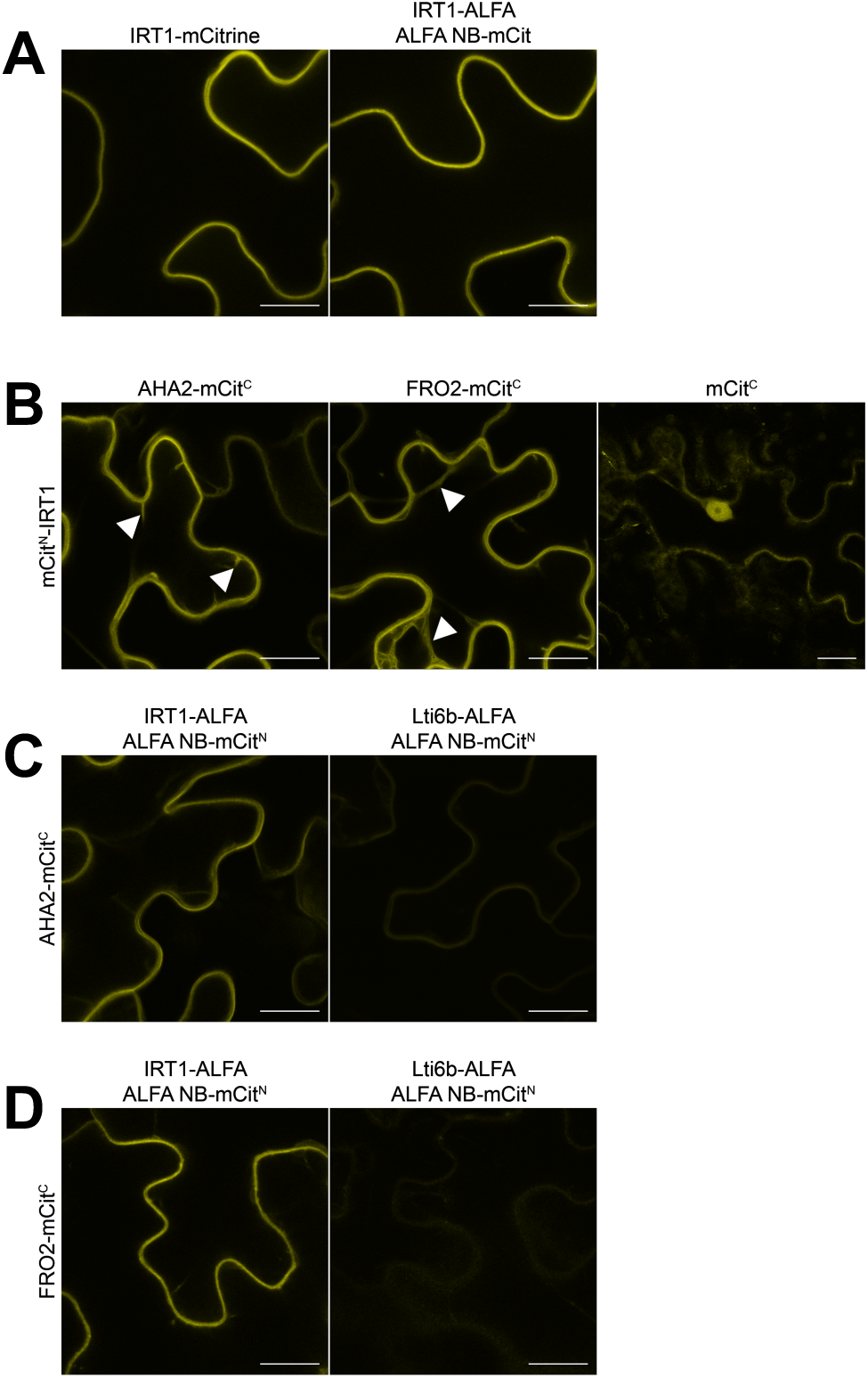
IRT1 Trimolecular Fluorescence Complementation (TriFC) assay. (A) Confocal microscope images of *N. benthamiana* transiently expressing IRT1-mCit or co-expressing IRT1-ALFA-mCit and ALFA NB-mCit. (B) Bimolecular Fluorescence Complementation (BiFC) assay with mCitN-IRT1 and, from left to right, AHA2-mCitC, FRO2-mCitC or mCitC alone. (C) Trimolecular Fluorescence Complementation (TriFC) assay with IRT1-ALFA/ALFA-NB-mCitN and AHA2-mCitC or Lti6b-ALFA/ALFA NB-mCitN and AHA2-mCitC. (D) TriFC assay with, IRT1-ALFA/ALFA-NB-mCitN and FRO2-mCitC or Lti6b-ALFA/ALFA NB-mCitN and FRO2-mCitC. Scale bars 10µm.

### ALFA tag-based induced-proximity approaches with IRT1

Induced proximity is a powerful strategy for the acute regulation of proteins through loss or gain of function. This innovative mechanism relies on proximity-inducing entities to bring proteins together to modulate the function of a target and downstream cellular processes. We therefore investigated if the ALFA tag/ALFA NB could be implemented for induced proximity strategies, focusing on protein degradation. To this purpose, we evaluated the impact of the artificial recruitment of IRT1-ALFA to an E3 ubiquitin ligase RING domain on IRT1 protein levels. The RING domain from the IDF1 E3 ligase (Shin et al., 2013), which is known to ubiquitinate IRT1, was first used as a test case, and its RING_dead_ mutant counterpart as negative control. We generated an IRT1-ALFA-mCit fusion to directly image the fate of IRT1-ALFA in the presence/absence of ALFA NB-RING. The IRT1-ALFA-mCit fusion was fully functional, as visualized by the complementation of the *irt1_crispr_* null mutant (Fig. 5A). Coexpression IRT1-ALFA-mCit with NB-RING in *N. benthamiana* leaves prevented the accumulation of IRT1-ALFA-mCit. The presence of the ALFA tag is necessary to drive IRT1-ALFA-mCit degradation as IRT1-mCit without ALFA accumulated at the plasma membrane in the presence or absence of NB-RING. (Fig. 5B-5D). Co-expression of the inactive RING_dead_ coupled to the ALFA NB was also ineffective in decreasing IRT1-ALFA-mCit levels (Fig. S4A, B). Besides, co-expression of IRT1-ALFA-mCit with a RING-tagBFP fusion devoid of ALFA NB did not trigger IRT1-ALFA degradation (Fig. S4 A, B), indicating that the sole overexpression of a RING domain is not sufficient to trigger IRT1-ALFA degradation. Altogether, these observations clearly show that degradation of IRT1-ALFA is specifically achieved through induced proximity with the ALFA NB-RING domain.

**Figure 5.**
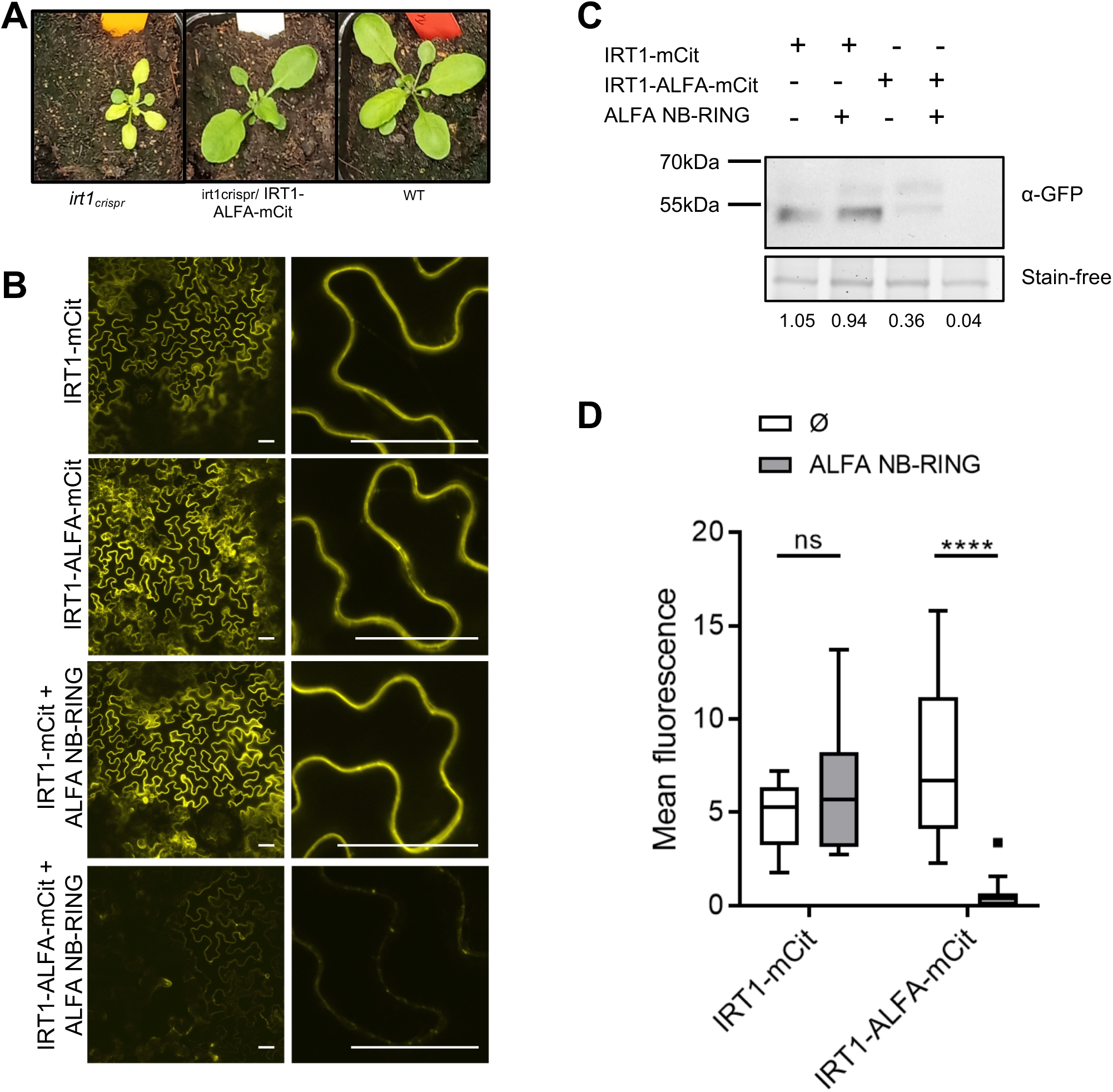
Targeted degradation of ALFA-tagged IRT1 protein. (A) Complementation of *irt1_crispr_* allele (Col-0) with IRT1-ALFA-mCit. Representative photos of *irt1_crispr_, irt1_crispr_*/IRT1-ALFA and wild-type plants are shown. (B) Confocal microscopy images of *N. benthamiana* expressing IRT1-mCit, IRT1-ALFA-mCit or co-expressing ALFA NB fused to RING domain of IDF1 (NB-RING) with IRT1-mCit or with IRT1-ALFA-mCit. Scale bar 50µm. (C) Western blot analysis of extracts from *N. benthamiana* transformation events presented in (B). The stain-free signal serves as a loading control. (D) Quantification of the total fluorescent signal of confocal microscopy images of *N. benthamiana* expressing the constructs shown in (B). Error bars represent standard deviation and asterisks indicate statistically significant differences [two-way ANOVA, Sidak post hoc test; ****P < 0.0001; ns, not significant].

### ALFA tag-based super-resolution microscopy

Protein tagging with an ALFA tag also allows for super-resolution microscopy analyses. This is especially interesting in the context of proteins reluctant to be tagging with large fluorescent proteins or with extracellular extremities where classical N- and C-terminal fusion with fluorescent protein suffer from extracellular acidic pH. To demonstrate the applicability of ALFA tagging to super-resolution microscopy, we performed single particle tracking Photoactivated Localization Microscopy (sptPALM) using the well-established Lti6b membrane protein that serves as a reference for sptPALM imaging of plasma membrane protein in plants (Hosy et al., 2015). To this purpose, we transiently expressed the Lti6b-mEos control or Lti6b-ALFA together with ALFA NB-mEos in *N. benthamiana* to follow the behavior of Lti6b at the cell surface. Both direct imaging of Lti6b using mEos or indirect imaging through Lti6b-ALFA/ALFA NB-mEos yielded comparable sptPALM profiles (Fig. 6A, B), mean squared displacements (Fig. 6D), distribution of the log of apparent diffusion coefficient (D) (Fig. 6E), and mean of peaks of apparent diffusion coefficient from thousand individual tracks (log (D)_Lti6b-mEos_= -1,02; log (D)_Lti6b-ALFA_= -1,00) (Fig 5F). In contrast, ALFA NB-mEos expressed alone showed irregular and scattered localization with relatively low density (Fig. S5A), and the apparent diffusion coefficient shows that the particles are divided into two populations, one with extremely short, immobile trajectories and the other highly mobile with longer trajectories, reflecting free diffusion in the cytosol (Fig. S5B-D). This argues for ALFA tagging as a suitable approach for super-resolution imaging of proteins otherwise not suited. To assess the potential of the ALFA NB approach for super-resolution imaging of proteins that are difficult to tag and with unfavorable topologies, we performed sptPALM imaging of IRT1. Contrary to the notorious mobility of the Lti6b that we confirmed here using both Lti6b-mEos or Lti6b-ALFA/ALFA NB-mEos, the behavior of IRT1-ALFA monitored with ALFA NB-mEos appeared radically different with more constrained diffusion (Fig. 6C-F) matching what has already been reported for other plasma membrane proteins such as AHA2 (Martinière et al., 2019). It is also worth noting that IRT1 has two particle populations, one more mobile and one immobile (log (D)_IRT1-ALFA immobile_= -2,20; log (D)_IRT1-ALFA mobile_= -1,65), reminiscent of what has already described for other proteins (Bademosi et al., 2018; Martinière et al., 2019).

**Figure 6.**
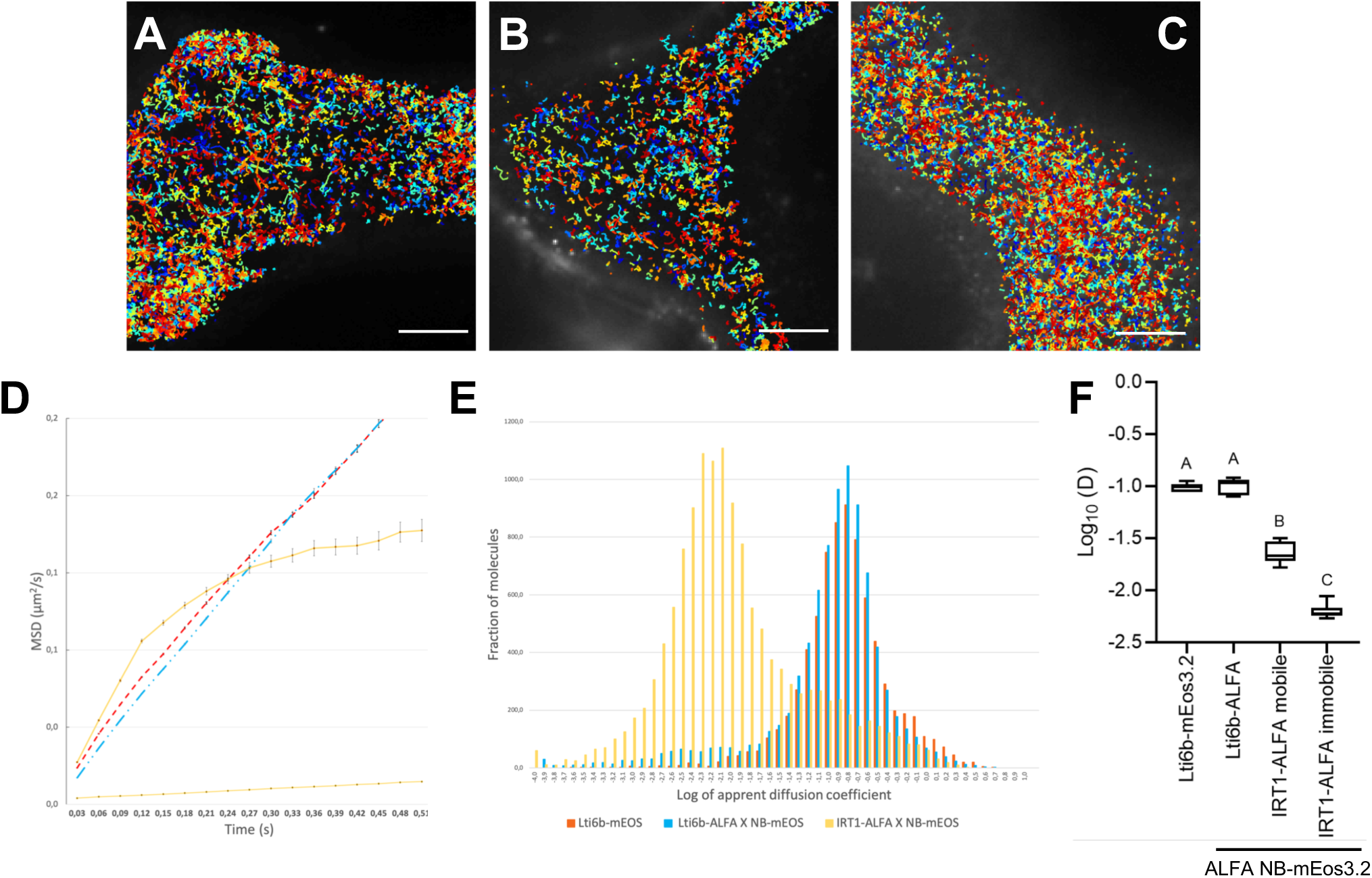
Super-resolution imaging of Lti6b-ALFA and IRT1-ALFA by sptPALM. (A-C) sptPALM trajectory map of a representative *N. benthamiana* cell expressing Lti6b-mEos3.2 (A), co-expressing Lti6b-ALFA and ALFA NB-mEos3.2 (B), co-expressing IRT1-ALFA and ALFA NB-mEos3.2 (C). Scale bars = 5µm. (D) Mean square displacement (MSD) over time for the global trajectories of Lti6b-mEOS3.2 (red), Lti6b-ALFA co-expressed with ALFA NB-mEOS3.2 (blue), IRT1-ALFA co-expressed with ALFA NB-mEOS3.2 mobile and immobile fractions (yellow). (E) Distribution of Log10 (Diffusion coefficient, D) of Lti6b-mEOS3.2 (red), Lti6b-ALFA co-expressed with ALFA NBmEOS3.2 (blue) and IRT1-ALFA co-expressed with ALFA NB-mEOS3.2 (yellow). (F) Boxplot of the peak log10 (D) values from seven to eight individual cells from 4 individual plants used in (E), with an average number of validated trajectories for MSD calculation after ROI selection for each cell of 12,879 for Lti6b-mEos, 7,449 for Lti6b NB-mEos, and 21,341 for IRT1 NB-mEos. Different letters indicate significant differences between conditions (one-way ANOVA, Tukey post hoc test, P < 0.001).

## Discussion

Protein tagging is essential in plant sciences to study protein function and regulation, as the availability of validated and currently commercialized antibodies targeting endogenous plant proteins remains limited. This is all the more true given that with emerging genome-editing approaches, one can consider knocking-in the tag at the endogenous locus to ensure faithful protein expression. Regardless, the nature of the tag is crucial to limit interference with protein folding and function, and for ease of detection to allow broad application. The ALFA tag is small, monovalent, devoid of lysine, hydrophilic without carrying any net charge, adopts a stable α-helical structure in solution, is absent within the proteome of relevant model organisms, and is recognized by the high affinity ALFA NB. The ALFA tag/ALFA NB technology has been successfully applied to the study of proteins in many organisms. In the present study, we have demonstrated that the ALFA tag and ALFA NB technology holds the premise to become the most versatile tag, allowing both high-end biochemical and imaging approaches in plants, and the preferred solution for when studying hard to tag proteins. Using the ALFA system allows the use of a single transgenic line or construct for transient expression harboring an ALFA-tagged target protein for a wealth of different applications including *in planta* applications. The broad range of applications of the ALFA technology provides the plant science community with a highly versatile tool, which will facilitate future scientific research. As designed, ALFA NB is genetically encoded and produced in the plant cell cytoplasm. Therefore, it cannot be used to recognize the ALFA tag when not facing the cytoplasm or the nucleus. We have however observed that binding of ALFA NB to membrane proteins can impact their delivery to the lytic vacuole by trapping them on the tonoplast. This phenomenon needs to be studied in greater detail, using electron microscopy approaches for example, to understand the endocytic sorting step at which ALFA NB interferes. But this observation also suggests that NB could be used to block target proteins in specific cellular compartments, notably to study the transfer to the vacuole or the recycling of membrane proteins. NBs could be an alternative to drugs, which can have pleiotropic effects that are sometimes difficult to analyze. The possibility to target the ALFA NB to various compartments using corresponding targeting signals (secretory pathway, extracellular space, organelles) will likely expand the use of the ALFA tag system to other biological questions. Besides, the breadth of application spans far beyond what we explored, with the possibility of fusing the NB to HRP for electron microscopy, TurboID for proximity labelling, post-translational modifiers (kinase, phosphatase, etc.) for targeted protein modification, or using the ALFA NB to target proteins *in vivo* to purify them and to increase the size of ALFA-tagged protein particles for cryoEM resolution of small proteins. In this study, we have shown the possibility of using the ALFA tag/ALFA NB system for targeted degradation approaches in a transient plant system with overexpression. These induced degradation strategies had already been used in various organisms, primarily using a GFP system (Caussinus et al., 2012; Wang et al., 2017) and very recently proof of concept using the ALFA tag and ALFA NB was achieved in animal cells (Yang et al., 2024). Further improvement of the ALFA technology for stable lines of plants for the study and understanding of cellular mechanisms will require finely controlled expression of the NB using inducible promoters or photocaged NB. ALFA-tag photobody where a photocaged ALFA NB variant cannot bind ALFA tag unless irradiated with light (365 nm) (Jedlitzke and Mootz, 2022). Light-induced ALFA photobody decaging will likely be a useful tool for optogenetic-based spatiotemporal control of proteins in plants.

The use of nanobodies fused with photoactivatable proteins for sptPALM approaches appears particularly promising, especially for membrane proteins with extracellular N- and C-termini, as most photoactivatable proteins used in super-resolution microscopy are highly sensitive to pH variations. By using a NB coupled to mEos, we were able to reproduce the results obtained with Lti6b directly fused to mEos, validating the use of this system for such approaches. We were also able to track single particles of the IRT1 transporter and assess its two diffusion profiles fairly confined within the plasma membrane of *N. benthamiana* cells. However, it will now be particularly interesting to use this approach in stable Arabidopsis lines and observe the distribution between these two diffusion profile changes of the IRT1 protein under metal stress conditions. Furthermore, to ensure the absence of background noise, which is indispensable for sptPALM, the use of destabilized nanobodies carrying a mutation of a few amino acids may turn out to be very handy. Such NB is degraded *in vivo* unless when bound to its target (Tang et al., 2016). Our sptPALM data showed a low background despite nanobody overexpression, likely due to the property of nanobodies to be naturally partially unstable when not attached to their antigen (Keller et al., 2018; Traenkle et al., 2020). The use of destabilized nanobodies would ensure the total absence of background for single-particle localization microscopy to facilitate particle tracking.

Overall, the non-exhaustive ALFA tag/ALFA NB toolkit presented here offers the most versatile tagging system for the characterization of proteins of interest in plants and will certainly be instrumental when dealing with hard to work proteins.

## Methods

### Constructs and transgenic plants

All the plasmids used in this study are listed in Table S1, and the primers used to clone the constructs are listed in Table S2. pDONR.P4P1R-UBI10 Gateway entry clones were previously described (Marquès-Bueno et al., 2016). The cDNAs or different targeting sequences were cloned into the gateway vectors pDONR221 and pDONR.P2RP3. The ALFA tag sequence optimized for *Arabidopsis* was added by PCR to cDNAs of interest or inserted into the PDONR221 and pDONR.P2RP3 gateway vectors. Vectors pDONR221-ALFA-IRT1-1 to 4 were cloned by inserting NotI restriction sites at different positions of pDONR221-IRT1 by PCR allowing the insertion of the ALFA tag by NotI digest followed by the ligation of the digested ALFA tag-NotI PCR fragment. The ALFA NB sequence optimized for *Arabidopsis* was synthesized and inserted into pDONR221. Final destination vectors combining the ALFA tag sequence and targeting sequences, or ALFA NB sequence and fluorescent proteins sequences for expression in plants, were assembled using the Gateway multisite recombination system (Life Technologies). Arabidopsis plants were transformed by floral dipping with the different constructs using *Agrobacterium tumefaciens* GV3101. To visualize NB relocalization, plants expressing constructs with different targeting sequences fused to the ALFA tag were crossed with plants expressing ALFA NB fused to other fluorescent proteins. Transgenic plants were selected on media containing the relevant antibiotic. To generate IRT1*_crispr_* lines, two guide RNAs (gRNAs) targeting IRT1 were cloned into plasmid pHEE401, as described in (Wang et al., 2015). The sgRNA sequences were selected using the CRISPR-P v2.0 online tool (Liu et al., 2017). T1 mutants were selected based on their chlorotic phenotype. Mutant plants were then backcrossed with wild-type Arabidopsis thaliana to eliminate the *Cas9* transgene.

### Plant growth conditions

Plants were grown vertically under sterile conditions at 21°C with a 16-hour light/8-hour dark cycle on half-strength Murashige and Skoog (MS/2) medium supplemented with 1% (w/v) sucrose and 1% (w/v) agar. For IRT1 localization experiments, plants were grown in the absence of iron and in the presence of physiological concentrations of IRT1 secondary substrates: Zn (15 µM), Mn (50 µM), and Co (0.05 µM) (-Fe +Metals) as described previously (Dubeaux et al., 2018).

### Transient Expression in Nicotiana benthamiana leaves

*Agrobacterium tumefaciens* GV3101 strain harboring the relevant constructs (including P19) were grown overnight and then resuspended in an infiltration solution containing 10 mM MgCl₂, 10 mM MES (pH 5.6), and 150 µM acetosyringone. Bacterial cultures were adjusted to a final OD₆₀₀ of 0.5 for each construct. The cultures were mixed and infiltrated into 4-week-old *N. benthamiana* leaves, and they were cultivated in a growth chamber for 48h before imaging.

### BiFC/TriFC Assay

To generate the final constructs for BiFC and TriFC experiments, pDONR221 or pDONR.P2RP3 plasmids carrying the N-terminal (mCitN) or C-terminal (mCitC) fragments of mCitrine (residues 1–155 and 155–238, respectively) were combined with pDONR221 or pDONR.P2RP3 plasmids carrying the protein of interest (POI) or ALFA NB, along with the pDONR.P1RP4-UBI10 promoter plasmid into a pDEST vector pK7m34GW or pH7m34GW^1^.

### Confocal microscopy

Confocal imaging was performed using a Leica SP8 confocal laser scanning microscope. mCit was excited at 514 nm and emission was collected from 525 to 550 nm, mSca was excited at 561 nm and emission was collected from 590 to 650 nm, and tagBFP was excited at 405 nm and emission was collected from 445 to 475 nm. For ALFA NB relocalization with control proteins, 8-day-old seedlings grown in half-strength LS medium were used. For ALFA NB relocalization with IRT1-ALFA, 10-day-old seedlings grown on half-strength Murashige and Skoog medium, then incubated in liquid medium containing either 50µM ferrozine or 50µM ferrozine plus an excess of non-iron metals Zn (300µM), Mn (1mM), Co (1µM) for 24 hours were used. Quantification of fluorescence was done as previously described (Spielmann et al., 2022).

### Western-blot analyses and immunoprecipitation

For Western blot analysis, total proteins were extracted from 100-300 mg of root of 15-day-old iron-deficient seedlings co-expressing IRT1-ALFA and ALFA NB-mCit in Laemmli buffer using a 3:1 ratio (w/v). Protein detection was performed using polyclonal anti-IRT1 (Agrisera AS11 1780; 1:5,000), horseradish peroxidase (HRP)-conjugated anti-GFP (Miletneyi; 1:5,000), and HRP-conjugated anti-rabbit (Bio-Rad; 1:20,000). Detection of HRP chemiluminescence was done using SuperSignal West Dura Extended Duration Substrate (Thermo Scientific) on a ChemiDoc MP Imaging System (Bio-Rad). IP was performed using 100-300 mg of root as previously described (Barberon et al., 2014; Dubeaux et al., 2018).

### Targeted Degradation Assay

For targeted degradation experiments of IRT1, vectors expressing IRT1-ALFA-mCitrine or IRT1-mCitrine were co-expressed in *N. benthamiana* alongside with either an ALFA NB fused to a functional or non-functional RING domain or mCherry fused to a C-terminal RING domain. 48 hours post-infiltration, leaves were analyzed using confocal microscopy or collected for Western blot analysis to observe IRT1 degradation.

### Single-Particle Tracking Photoactivated Localization Microscopy (sptPALM) analysis

For sptPALM analysis, plasmids expressing ALFA-POI and NB-mEos3.2 were transformed in Agrobacterium, and co-infiltrated into tobacco leaves 48h before imaging. Small leaf pieces were mounted on a drop of 1.8% agarose in 1X PBS between the slide and coverslips using a Gene Frame^®^ sealing system (Thermo Scientific).

SMLM (Single-molecule localization microscopy) was performed with an automated inverted epifluorescence microscope Nikon Ti-E/B equipped with the “perfect focus system” (PFS, Nikon), a CFI Apochromat TIRF 100XC Oil Objectif (NA1.49), a quad-band dichroic mirror (97335 Nikon N-STORM TIRF Filter Set), a TIRF illumination arm (here we use a highly inclined and laminated optical sheet HILO illumination mode), an EM-CCD ANDOR iXon Ultra DU897 camera (Pixel Size 171 nm) and a thermostatic chamber at 24°C to limit the instrumental drift. Excitation is controlled with an AOTF. For spt-PALM, we used to control mEos photoconversion a 405nm laser (CoherentTM - Cube 405-100C) at low laser power and for imaging a 561nm laser (CoherentTM – Sapphire) used at a power of ∼0.1kW/cm². For spt-PALM image sequence acquisition, imaging laser (561 nm) was used in continuous and acquisition time was set at 30ms. Images were captured using Nis-Elements AR software (Nikon).

The images obtained were analyzed using the workflow described previously (Bayle et al., 2021). In summary, the 5,000 image stacks obtained are divided into 5 stacks of 1,000 images by running the macro ‘Stack_split_macro.ijm’. These files were then loaded into the MTT software keeping the default parameters, with data output in .mat and alpha threshold at 1000. To analyze the trajectories obtained and compile the data, sptPALM-viewer was used. After selecting the ROI, the following parameters were used: 3 maximum number of blinks, 7 frames for minimum length trajectory, 0,50 maximum display time for the MSD, and 4 points to calculate apparent D.

## Acknowledgements

We would like to thank the FRAIB and the LITC imaging platforms of Toulouse TRI-Genotoul for their assistance. This work was supported by research grants from the French National Research Agency (ANR-21-CE20-0046 to G.V. and 22-PESV-0002 to J.N.) and the French Laboratory of Excellence (project “TULIP” grant nos. ANR–10–LABX–41 and ANR–11– IDEX–0002–02 to G.V.).

