## Supplemental Figures and Tables for "Broad application of plant protein tagging with ALFA tag for nanobody-based imaging and biochemical approaches"

[3] Co-first author

### Supplemental figure legends

**Figure S1. Expression of ALFA-NB-Fluorescent protein *in planta.*** (A) Schematic illustration of ALFA NB fused to FP expressed in Arabidopsis. (B-G) Confocal microscopy images of Arabidopsis plants expressing ALFA NB-tagBFP in leaves (B), roots (C), root tips (D), or to NB-mSca in leaves (E), roots (F), and root tips (G). Scale bar 50µm. (H) Statistical analysis of root length from 14-day-old Arabidopsis wild-type and transgenic seedlings expressing different ALFA NB-FP on half-strength MS medium. Experiments were carried out in triplicates. No statistical difference was observed (one-way ANOVA, Sidak post hoc test; ns, not significant). (I) The phenotype of wild-type (Col-0) and ALFA NB-FP-expressing plants.

**Figure S2. Detection of ALFA-tagged MAP4 by ALFA NB fused to fluorescent proteins**. (A-B) Confocal microscopy images of hypocotyls from 10-day-old seedlings of *A. thaliana* co-expressing MAP4-ALFA and ALFA NB-mCit before oryzalin treatment (A), or after a 2h oryzalin treatment (B). Scale bars 5µm.

**Figure S3. Detection of ALFA-tagged reporter proteins by ALFA NB fused to fluorescent proteins.** (A-B) Confocal microscopy images of *A. thaliana* co-expressing the mitochondrial inner-membrane (IM) signal fused to mCit-ALFA and ALFA NB-mSca (A), and the endoplasmic reticulum HDEL-mSca-ALFA and ALFA NB-mCit (B). Scale bars 10µm.

**Figure S4. Targeted degradation of IRT1 protein.** (A) Confocal microscopy images of *N. benthamiana* transiently co-expressing IRT1-ALFA-mCit with ALFA-NB fused to the RING domain of IDF1 (NB-RING), ALFA-NB fused to the inactive RING domain of IDF1 (NB-

RING$_{dead}$) or with tag-BFP fused to the RING domain of IDF1 (BFP-RING). Scale bar 50µm. Scale bar 50µm. (B) Quantification of the total fluorescent signal of confocal microscopy images of *N. benthamiana* expressing the constructs shown in (B). Error bars represent standard deviation and asterisks indicate statistically significant differences [two-way ANOVA, Sidak post hoc test ; ****P < 0.0001; ns, not significant] .

**Figure S5. sptPALM of ALFA-tagged proteins.** (A) sptPALM trajectory map of a representative *N. benthamiana* cell expressing ALFA NB-mEos3.2. Scale bar=5 µm. (B) Mean square displacement (MSD) over time with ALFA NB-mEos3.2 as a control added to the graphical data in Figure 5. (C) Distribution of Log10 (Diffusion coefficient, D) with NB-mEos3.2 (green) as a control added to the graphical data in Figure 5. (D) Boxplot of the peak log10 (D) values from eight different cells from 4 individual plants used in (C) with an average number of validated trajectories for MSD calculation after ROI selection for each cell of 4743 for ALFA NB-mEos3.2 mobile and immobile fractions as a control added to the graphical data in Figure 5. Different letters indicate significant differences between conditions (one-way ANOVA, Tukey post hoc test, P < 0.001).

**Figure S6. Uncropped western blot.** (A) Uncropped western blot of Figure 3. (B) Uncropped western blot of Figure 5.

**Figure S1.**

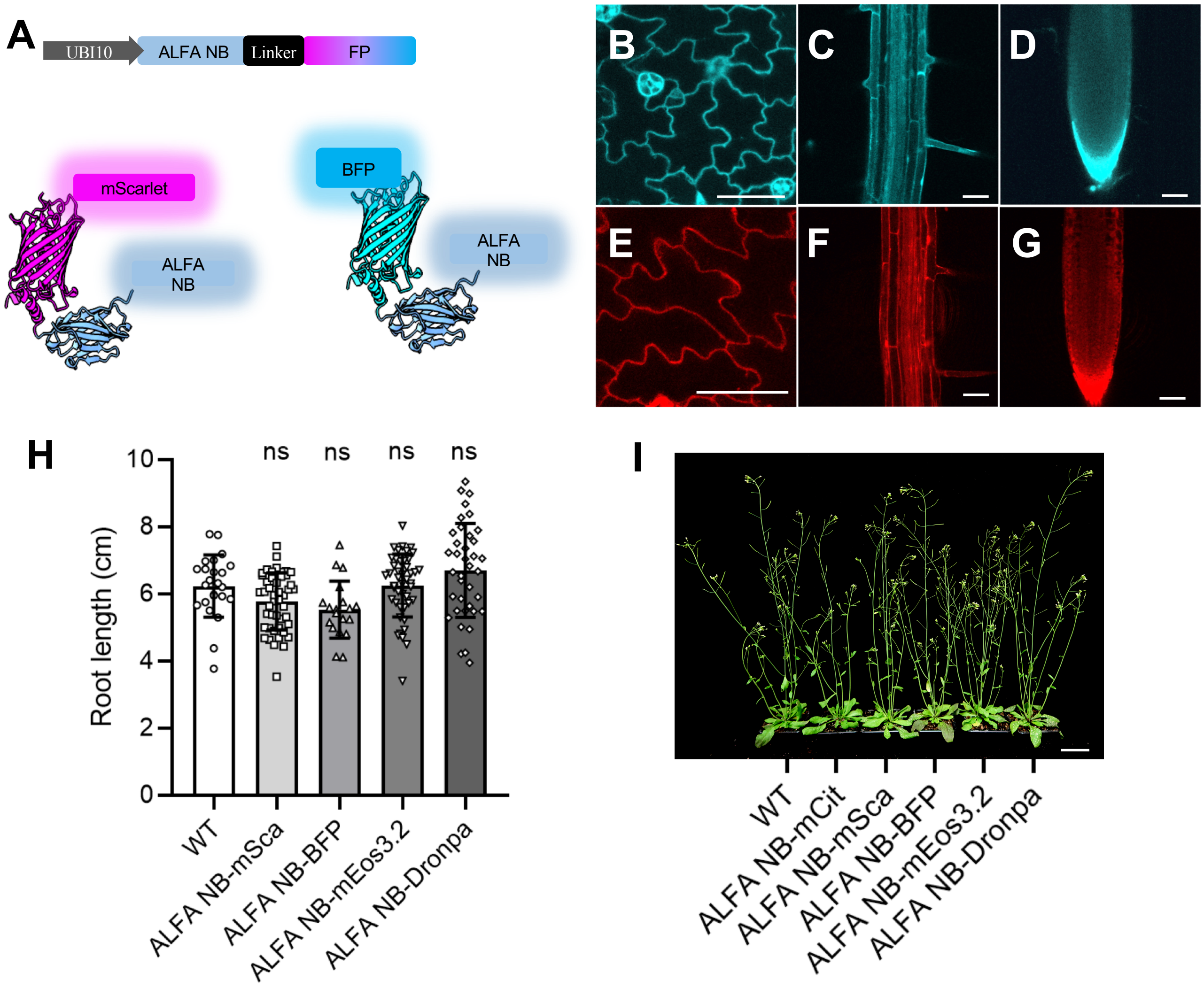
A
UBI10
ALFA NB
Linker
FP
mScarlet
ALFA NB
BFP
ALFA NB
B
C
D
E
F
G
H
Root length (cm)
ns
ns
ns
ns
WT
ALFA NB-mSca
ALFA NB-BFP
ALFA NB-mEos3.2
ALFA NB-Dronpa
I
WT
ALFA NB-mCit
ALFA NB-mSca
ALFA NB-BFP
ALFA NB-mEos3.2
ALFA NB-Dronpa

**Figure S2.**

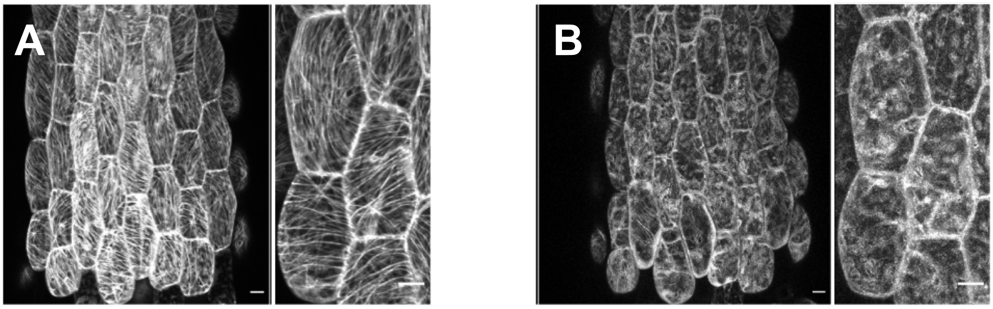

### Figure S3.

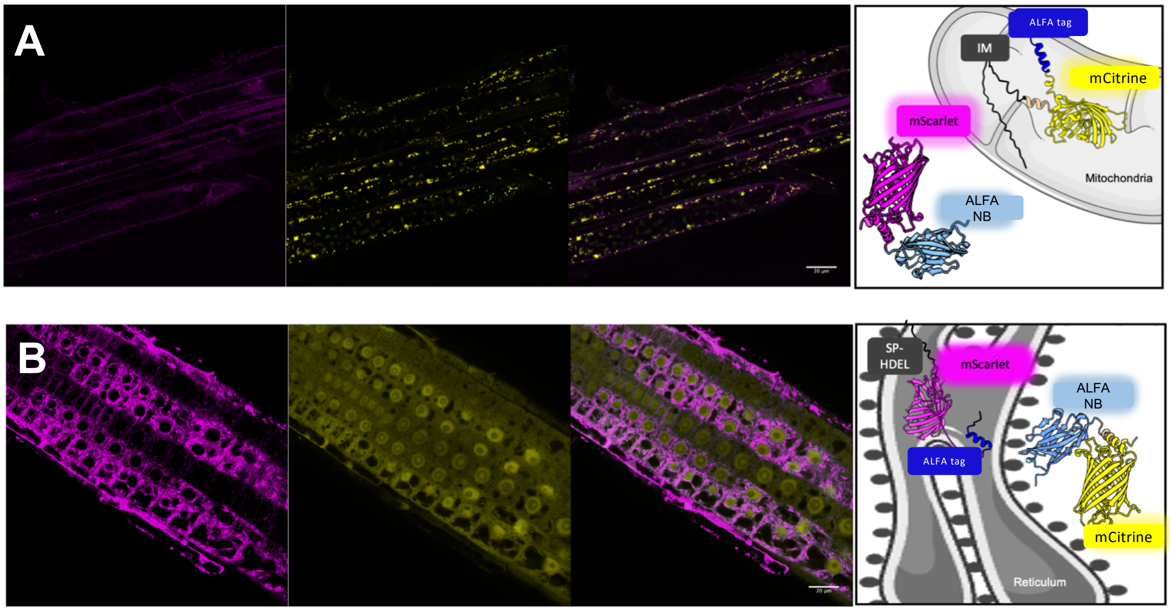
A
ALFA tag
IM
mCitrine
mScarlet
Mitochondria
ALFA
NB
B
SP-
HDEL
mScarlet
ALFA
NB
ALFA tag
mCitrine
Reticulum

**Figure S4.**

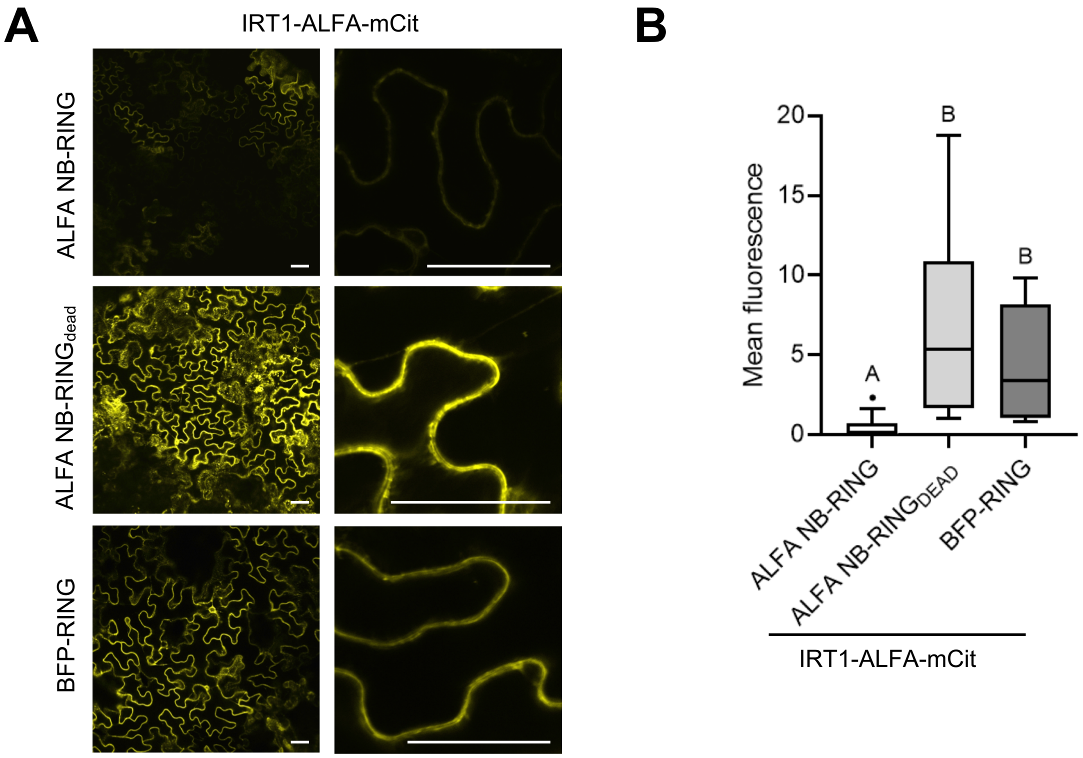
A
IRT1-ALFA-mCit
ALFA NB-RING
ALFA NB-RINGdead
BFP-RING
B
Mean fluorescence
20
15
10
5
0
A
B
B
ALFA NB-RING
ALFA NB-RINGDEAD
BFP-RING
IRT1-ALFA-mCit

**Figure S5.**

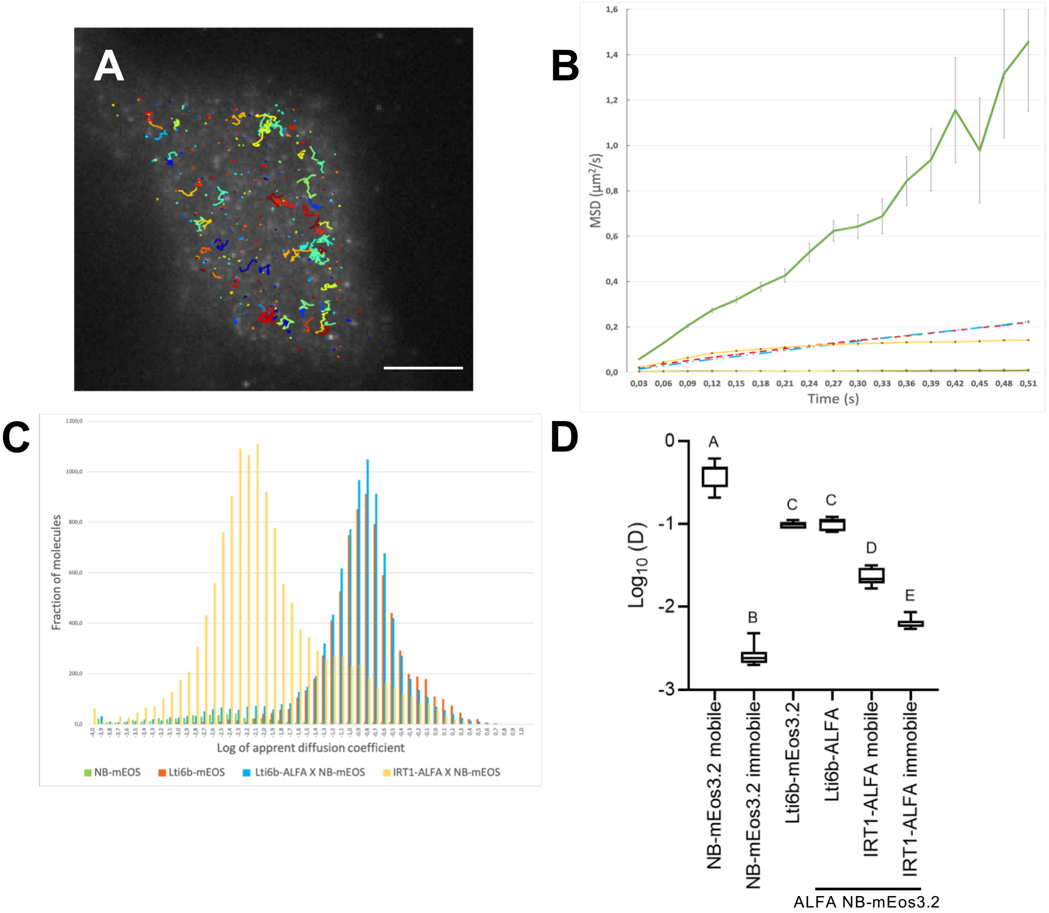

A
B
MSD (µm²/s)
Time (s)
0,03 0,06 0,09 0,12 0,15 0,18 0,21 0,24 0,27 0,30 0,33 0,36 0,39 0,42 0,45 0,48 0,51
0,0 0,2 0,4 0,6 0,8 1,0 1,2 1,4 1,6
C
Fraction of molecules
Log of apprent diffusion coefficient
NB-mEOS
Lti6b-mEOS
Lti6b-ALFA X NB-mEOS
IRT1-ALFA X NB-mEOS
D
Log10 (D)
0
-1
-2
-3
A
B
C
C
D
E
NB-mEos3.2 mobile
NB-mEos3.2 immobile
Lti6b-mEos3.2
Lti6b-ALFA
IRT1-ALFA mobile
IRT1-ALFA immobile
ALFA NB-mEos3.2

**Figure S6.**

**A**

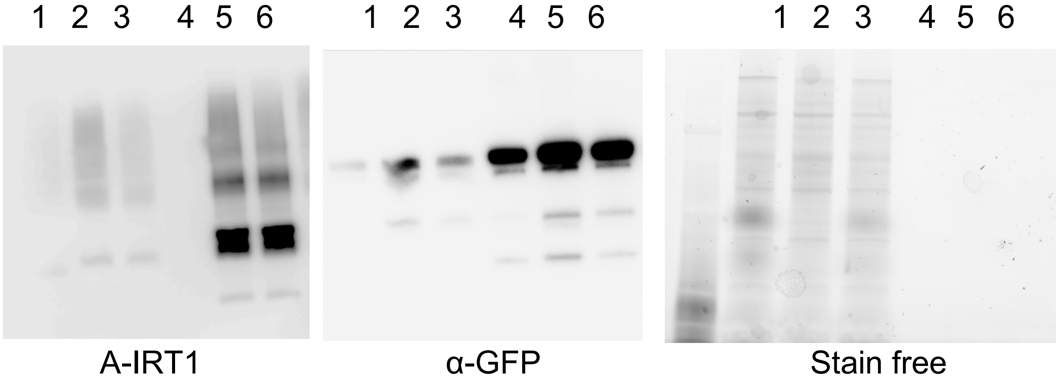


1 = ALFA NB-mCit INPUT
2 = ALFA NB-mCit + IRT1-ALFA INPUT
3 = ALFA NB-mCit + IRT1-ALFA INPUT
4 = ALFA NB-mCit IP
5 = ALFA NB-mCit + IRT1-ALFA IP
6 = ALFA NB-mCit + IRT1-ALFA IP

**B**

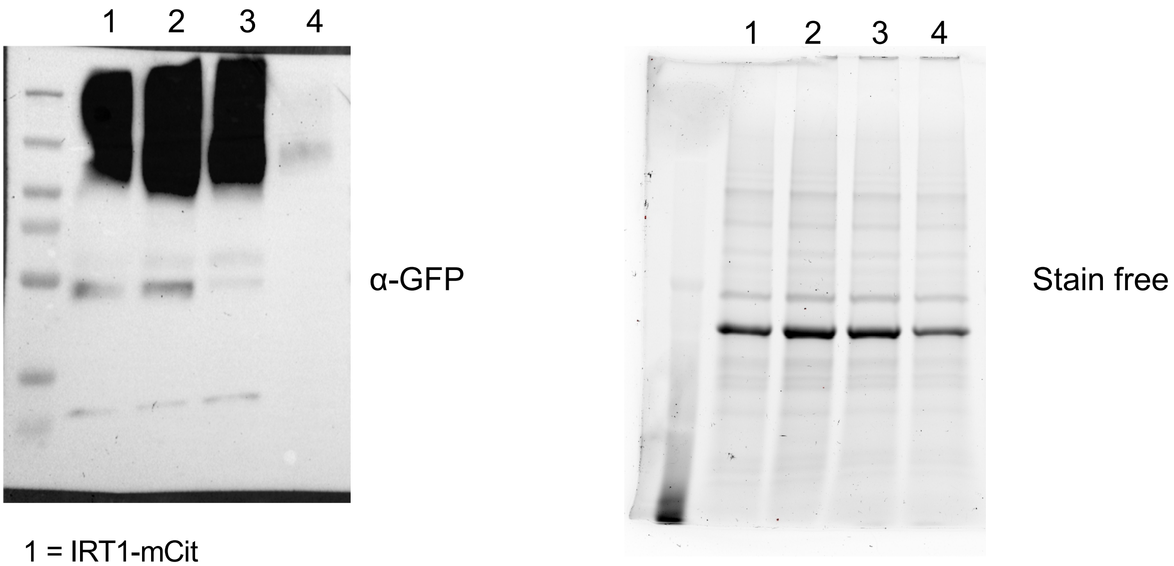


1 = IRT1-mCit
2 = IRT1-mCit + ALFA NB-RING
3 = IRT1-ALFA-mCit
4 = IRT1-ALFA-mCit + ALFA NB-RING

**Table 1.** Liste of constructs used or generated in this study**.**

| plasmids | origin |
|---|---|
| PDONOR221 ALFA-tag | this paper |
| P2RP3 ALFA-tag | this paper |
| PDONOR221-IRT1 ALFA-tag position1 | this paper |
| PDONOR221-IRT1 ALFA-tag position 2 | this paper |
| PDONOR221-IRT1 ALFA-tag position 3 | this paper |
| PDONOR221-IRT1 ALFA-tag position 4 | this paper |
| P2RP3 MAP4 | this paper |
| PDONOR221 SP-mScarlet-HDEL | this paper |
| P2RP3 ALFA-tag HDEL | this paper |
| PDONOR221-NLS-mCitrine | this paper |
| P2RP3-Lti6b w STP | this paper |
| PDONOR221 OM64 | this paper |
| PDONOR221 IM mito-DRONPA | this paper |
| PDONOR221 mCitrine ALFAtag | this paper |
| PDONOR221 mScarlet ALFAtag | this paper |
| P2RP3 mCitrine ALFAtag | this paper |
| P2RP3 mScarlet ALFAtag | this paper |
| pDONOR221 ALFA-Nanobody | this paper |
| P2RP3 mCitrine | Dubeaux et al. 2018 |
| P2RP3 mScarlet | this paper |
| P2RP3 BFP | This paper |
| P2RP3 DRONPA | this paper |
| P2RP3 PACherry | this paper |
| P1RP4 promoUBI10 | Marquès-Bueno et al., 2016 |
| P1RP4 promoIRT1 | Marquès-Bueno et al., 2016 |
| pG7m34GW pIRT1:ALFAtag -IRT1 position 1 | this paper |
| pG7m34GW pIRT1:ALFAtag -IRT1 position 2 | this paper |
| pG7m34GW pIRT1:ALFAtag -IRT1 position 3 | this paper |
| pG7m34GW pIRT1:ALFAtag -IRT1 position 4 | this paper |
| pK7m34GW pUBI10:ALFAtag -Lti6b | this paper |
| pG7m34GW pUBI10:mEOS -Lti6b | this paper |
| pK7m34GW pUBI10:OM64-ALFA-tag | this paper |
| pH7m34GW pUBI10:mScarlet-HDEL-ALFAtag | this paper |
| pK7m34GW pUBI10:ALFAtag-MAP4 | this paper |
| pG7m34GW pUBI10:NLS-mCitrine-ALFAtag | this paper |
| pK7m34GW pUBI10:IN-MITO-DRONPA-ALFAtag | this paper |
| pG7m34GW pUBI10:ALFAtag-mCitrine-Lti6b | this paper |
| pK7m34GW pUBI10:ALFAtag-mCitrine-MAP4 | this paper |
| pG7m34GW pUBI10:ALFA Nanobody-mCitrine | this paper |
| pG7m34GW pUBI10:ALFA Nanobody-mScarlet | this paper |
| pG7m34GW pUBI10:ALFA Nanobody-tagBFP | this paper |
| pG7m34GW pUBI10:ALFA Nanobody-DRONPA | this paper |
| pB7m34GW 2X35S:Alfa-nanobody:mCit$^{N}$ | this paper |
| pH7m34GW 2X35S:AscI-AHA2-PacI:mCit$^{C}$ | this paper |
| pH7m34GW 2X35S:AscI-FRO2-PacI:mCit$^{C}$ | this paper |

**Table 2.** Liste of primers used in this study**.**

| Name | Sequence 5' to 3' | Purpose |
|---|---|---|
| Not-IRT1-1 fw | taattGCGGCCGCggcggatcccgggttcgctgac | Subcloning NotI restriction site in position 1 of PDONOR221 IRT1 |
| Not-IRT1-1 rev | ttataGCGGCCGCagccggatcctgcgtcaaca | Subcloning NotI restriction site in position 1 of PDONOR221 IRT1 |
| Not-IRT1-2 fw | TAAAGCggccGCGTCGACGAGTACTcgtaacgtttcgttcctccaa | Subcloning NotI restriction site in position 2 of PDONOR221 IRT1 |
| Not-IRT1-2 fw | TATAGCggccGCGTTTAAACCTGCAGCgctaaagagaggagctccaa | Subcloning NotI restriction site in position 2 of PDONOR221 IRT1 |
| Not-IRT1-3 fw | TAAAGCGGCCGCGTCGACGAGTACTagc cta tac acc agc aag aac gca gt | Subcloning NotI restriction site in position 3 of PDONOR221 IRT1 |
| Not-IRT1-3 Rv | TATAGCggccGCGTTTAAACCTGCAGCcgtggccatggagtcaatggcta | Subcloning NotI restriction site in position 3 of PDONOR221 IRT1 |
| Not-IRT1-4 fw | TAAAGCggccGCGTCGACGAGTACTatc aaa atg cag ttc aag tgt | Subcloning NotI restriction site in position 4 of PDONOR221 IRT1 |
| Not-IRT1-4 rev | TATAGCggccGCGTTTAAACCTGCAGCgctaccttgaagcttaggtcccat | Subcloning NotI restriction site in position 4 of PDONOR221 IRT1 |
| IRT1-ALFA-f | TCCTTTTAACAGCGATCGCGTATTTCGTCT | Cloning PDONR IRT1 PCR fragment to make PDONOR221-IRT1 ALFA-tag positions 1, 2, 3 and 4 |
| IRT1-ALFA-r | AGACGAAATACGCGATCGCTGTTAAAAGGA | Cloning PDONR IRT1 PCR fragment to make PDONOR221-IRT1 ALFA-tag positions 1, 2, 3 and 4 |
| ALFAtag | GT TTA AAC GCG GCC GCGccttctagacttgaggaagagct | Cloning ALFAtag-NotI PCR fragment to make PDONOR221-IRT1 ALFA-tag positions 1, 2, 3 and 4 |
| ALFAtag rev | TCGACGCGGCCGCaggttcagtaagtctccttcga | Cloning ALFAtag-NotI PCR fragment to make PDONOR221-IRT1 ALFA-tag positions 1, 2, 3 and 4 |
| SP-mscarlet fw | GGGG ACA AGT TTG TAC AAA AAA GCA GGC Tgaaccatgaagactaatctttttctct | Cloning SP-mSarlet-HDEL PCR fragment to make Cloning PDONR221 SP-mScarlet |
| SP-mScarlet rev | GGGGACCACTTTGTACAAGAAAGCTGGGTccttgtacagctcgtccatgc | Cloning SP-mSarlet-HDEL PCR fragment to make Cloning PDONR221 SP-mScarlet |
| ALFA-HDEL-Fw | aagctcatcatgaggttcagtaagtctccttcga | Cloning ALFA-HDEL PCR fragment ALFA-tag HDEL to make P2RP3 ALFAtag-HDEL |
| ALFA-HDEL-rev | GGGGACAACTTTGTATAaTAAAGTTGCttaaagctcatcatgagg | Cloning ALFA-HDEL PCR fragment ALFA-tag HDEL to make P2RP3 ALFAtag-HDEL |
| OM64 Fw | GGGG ACA AGT TTG TAC AAA AAA GCA GGC TgaaccATGTCGAATACGCTTTCTTTGATTC | Cloning OM64 PCR fragment to make pDONR221 OM64 |
| OM64 RV | GGGGACCACTTTGTACAAGAAAGCTGGGTcAGGGAAAGGAAGAAGCTCGAAACG | Cloning OM64 PCR fragment to make pDONR221 OM64 |
| attB1 NLS | GGGG ACA AGT TTG TAC AAA AAA GCA GGC Tgaacc ATGCCAAAGAAGAAGAGGAAGGTT | Cloning NLS-mCitrine PCR fragment to make pDONOR221-NLS-mCitrine |
| attB1 IM mito Fw | GGGG ACA AGT TTG TAC AAA AAA GCA GGC Tga acc ATGCTTTCACTACGTCAATCT | Cloning MITO PCR fragment to make PDONR221 MITO |
| attB2 IM mito Rv | GGGG AC CAC TTT GTA CAA GAA AGC TGG GTc GGTGGCGACCGGTGgatcCCGT | Cloning MITO-Dronpa PCR fragment to make PDONR221 MITO-Dronpa |
| attB2 DRONPA Rv | GGGG AC CAC TTT GTA CAA GAA AGC TGG GTc CTTGGCCTGCCTCGGCA | Cloning MITO-Dronpa PCR fragment L to make PDONR221 MITO-Dronpa |
| attB ALFA fw | GGGG ACA AGT TTG TAC AAA AAA GCA GGC Tgaacc ATG ccttctagacttgaggaagagct | Cloning ALFA-tag PCR fragment to make PDONR221 ALFA-tag |
| ALFA rev | *tgaggaagagcttcgaaggagacttactgaacctgccggaggcggtggat* | Cloning ALFA-tag PCR fragment to make PDONR221 ALFA-tag |
| attB ALFA rev | GGGGACCACTTTGTACAAGAAAGCTGGGT*catccaccgcctccggc* | Cloning ALFA-tag PCR fragment to make PDONR221 ALFA-tag |
| pDONR fw IRT1 | GT TTA AAC GCG GCC GCGccttctagacttgaggaagagct | Cloning IRT1pDONR PCR fragment to make PDONR221 IRT1 ALFA-tag |
| pDONR rv IRT1 | TCGACGCGGCCGCaggttcagtaagtctccttcga | Cloning IRT1pDONR PCR fragment to make PDONR221 IRT1 ALFA-tag |
| attB2R ALFA | GGGGACAGCTTTCTTGTACAAAGTGGCA ccttctagacttgaggaaga | Cloning ALFA-tag PCR fragment to make P2RP3 ALFA-tag |
| attB3 ALFA | GGGGACAACTTTGTATAaTAAAGTTGCttaaggttcagtaagtctcct | Cloning ALFA-tag PCR fragment to make P2RP3 ALFA-tag |
| AscI-AHA2-Fwd | GGGGCGCGCCATGTCGAGTCTCGAAGATAT | Cloning AHA2 PCR fragment to make pH7m34GW 2X35S:AscI-AHA2-PacI:mCit$^{C}$ |
| PacI-AHA2-Rev | GGTTAATTAACACAGTGTAGTGACTGGGAG | Cloning AHA2 PCR fragment to make pH7m34GW 2X35S:AscI-AHA2-PacI:mCit$^{C}$ |
| AscI-FRO2-Fwd | GGGGCGCGCCATGGAGATCGAAAAAAGCAATA | Cloning FRO2 PCR fragment to make pH7m34GW 2X35S:AscI-FRO2-PacI:mCit$^{C}$ |
| PacI-FRO2-Rev | GGTTAATTAACCAGCTGAAACTGATAGATTC | Cloning AHA2 PCR fragment to make pH7m34GW 2X35S:AscI-AHA2-PacI:mCit$^{C}$ |
